# N-terminal phosphorylation inhibits Arabidopsis katanin and affects vegetative and reproductive development in opposite ways

**DOI:** 10.64898/2026.01.20.700631

**Authors:** Vivek Ambastha, Graham Burkart, Rachappa Balkunde, Ram Dixit

**Affiliations:** Department of Biology, Washington University in St. Louis, St. Louis, MO 63130, USA

## Abstract

Katanin is an evolutionarily conserved microtubule-severing enzyme that is essential for cytoskeletal remodeling throughout the plant life cycle. However, the molecular mechanisms that tune katanin activity to meet distinct cellular requirements remain unclear. Here, we demonstrate that N-terminal phosphorylation of the *Arabidopsis thaliana* p60 katanin subunit (KTN1) serves as a key regulatory switch controlling microtubule severing during vegetative and reproductive development. Using *in vitro* biochemical assays, we show that combined phosphorylation of three conserved serine residues (S92, S147, S199) inhibits KTN1’s microtubule-severing activity by reducing both microtubule-binding affinity and ATPase activity. Strikingly, phosphomimetic (DDD) and phosphonull (AAA) versions of KTN1 exhibit opposite developmental phenotypes. The constitutively active AAA mutant rescues defects in cortical microtubule organization and vegetative growth but leads to abnormal meiotic spindles, reduced pollen viability, and defective pollen tube growth, resulting in low male fertility. Conversely, the catalytically impaired DDD mutant fails to restore vegetative growth but supports normal male fertility. These findings reveal that phosphorylation differentially modulates KTN1 activity to balance the opposing requirements for high microtubule severing during interphase cell expansion versus limited severing during meiotic cell divisions, providing a sophisticated mechanism to coordinate cytoskeletal dynamics with plant developmental programs.

## INTRODUCTION

Microtubules are dynamic cytoskeletal polymers underlying essential cellular processes in plants, including cell division, morphogenesis, and intracellular transport. During interphase, cortical microtubules align beneath the plasma membrane and direct the deposition of cellulose microfibrils and other cell wall components (Paredez et al. 2006; McFarlane et al. 2008; Kong et al. 2015; Zhu et al. 2015), thereby controlling the axis of cell expansion. Cortical microtubules form characteristic arrays in different cell types and can rapidly change their pattern in response to developmental and environmental signals (Hamant et al. 2008; Zhang et al. 2011; Lindeboom et al. 2013; Sampathkumar et al. 2014). During mitosis, microtubules reorganize into specialized arrays including the preprophase band, spindle apparatus, and phragmoplast, which function to specify the cell division plane, segregate chromosomes, and orchestrate cytokinesis, respectively (Motta and Schnittger 2021). Similar microtubule reorganization occurs during meiosis to mediate two rounds of cell division to produce four haploid gametes (Prusicki et al. 2019).

Remodeling of the microtubule cytoskeleton, which underpins its functional versatility, is mediated by microtubule-associated proteins which control microtubule nucleation, dynamics, and higher-order assembly.

Among the microtubule-associated proteins that regulate microtubule dynamics, katanin has emerged as a critical factor for the construction and remodeling of microtubule arrays in both plants and animals. Katanin is a heterodimeric complex composed of a catalytic p60 subunit containing an AAA ATPase domain and a regulatory p80 subunit (McNally and Vale 1993; Hartman et al. 1998). The p60 catalytic subunit is sufficient for microtubule severing activity and functions by forming ATP-dependent hexamers that encircle the electronegative C-terminal tails of tubulin subunits projecting from the microtubule surface (Hartman et al. 1998; Zehr et al. 2020). Through a mechanism involving conformational changes driven by ATP hydrolysis, katanin generates mechanical force to dislodge tubulin dimers from the microtubule lattice, ultimately leading to microtubule breakage (Zehr et al. 2017). The p80 regulatory subunit localizes the p60 subunit to specific subcellular sites and enhances microtubule-severing activity (Hartman et al. 1998; McNally et al. 2000).

The central role of katanin during plant growth and development is evident from the pleiotropic phenotypes observed in loss-of-function mutants (Bichet et al. 2001; Burk et al. 2001; Bouquin et al. 2003; Komorisono et al. 2005; Martinez et al. 2025). These mutants display disorganized interphase cortical microtubule arrays due to compromised microtubule severing at nucleation sites and crossover points where microtubules intersect (Lindeboom et al. 2013; Zhang et al. 2013). During cell division, katanin mutants exhibit aberrant preprophase bands consisting of poorly aligned microtubules, multipolar spindles, and phragmoplasts with elongated and bent microtubules (Panteris et al. 2011; Komis et al. 2017). Together, these microtubule abnormalities manifest as reduced anisotropic cell expansion, mispositioned cell division planes, organs of small size and abnormal shape, and reduced fertility.

The profusion of katanin functions is suggestive of the existence of multiple regulatory mechanisms to control the location, timing, and amount of katanin activity. The regulatory p80 subunit targets katanin to centrosomes and spindle poles in animal cells (Hartman et al. 1998; McNally et al. 2000), and to cortical microtubule nucleation and crossover sites in plants (Wang et al. 2017). Additional targeting factors have been identified in plants, including the mitosis-specific CORTICAL MICROTUBULE DISORDERING (CORD) proteins that recruit katanin to the distal phragmoplast zone to promote phragmoplast expansion through localized severing (Sasaki et al. 2019). Katanin activity is also regulated by microtubule-associated proteins that limit enzyme accessibility to the microtubule lattice. For example, the microtubule-bundling protein MAP65-1 inhibits katanin binding along microtubule sidewalls through lateral cross-linking of adjacent microtubules (Burkart and Dixit 2019). In animals, post-translational modifications of tubulin provide another layer of control by modulating katanin binding affinity and severing efficiency (Szczesna et al. 2022).

Accumulating evidence from animal systems indicates that post-translational modifications of p60 katanin itself, particularly ubiquitylation and phosphorylation, represent key regulatory mechanisms for controlling the amount and activity of this enzyme. In *Caenorhabditis elegans*, the p60 katanin homolog MEI-1 is essential for meiotic spindle assembly but must be rapidly inactivated before the first mitotic division to prevent spindle defects and embryonic lethality. This developmental switch is accomplished by the minibrain kinase MBK-2 which phosphorylates MEI-1 at multiple serine residues to inhibit ATPase activity and target MEI-1 for proteasomal degradation (Joly et al. 2020). During meiosis, protein phosphatases counteract the inhibitory phosphorylation to maintain katanin activity (Han et al. 2009; Gomes et al. 2013).

Similarly, in mammalian cells, phosphorylation of p60 katanin by the dual specificity tyrosine-regulated kinase 2 targets the enzyme for proteasomal degradation (Maddika and Chen 2009).

Studies in Xenopus egg extracts have revealed that phosphorylation can also directly inhibit p60 katanin’s catalytic activity without affecting protein turnover. Phosphorylation of *Xenopus laevis* p60 katanin at serine 131 by Aurora B kinase inhibits microtubule severing activity and leads to longer meiotic spindles compared to *X. tropicalis* which lacks the inhibitory phosphorylation site in its p60 subunit (Loughlin et al. 2011). Biochemical experiments revealed that phosphomimetic modification of serine 131 does not affect basal ATP hydrolysis or microtubule binding affinity but rather suppresses concentration-dependent oligomerization of p60 katanin on the microtubule surface (Whitehead et al. 2013). Thus, at high local concentrations phosphomimetic p60 can hexamerize and restore wild-type levels of severing activity, while the bulk cellular pool remains inhibited. Together, these findings establish that phosphorylation and dephosphorylation cycles can control katanin abundance and activity to provide the correct level of microtubule severing at specific locations and developmental stages.

Recent work in *Arabidopsis thaliana* has identified the conserved protein phosphatase PP2A as a katanin-interacting protein that dephosphorylates p60 katanin to promote cortical microtubule organization in petal conical cells (Ren et al. 2022), suggesting that phosphoregulatory mechanisms operate in plants as well. However, fundamental questions remain regarding how phosphorylation regulates katanin in plants and whether this regulatory mechanism contributes to the distinct microtubule-severing requirements during interphase versus mitotic and meiotic cell divisions. Here, we investigate the role of N-terminal phosphorylation in controlling Arabidopsis katanin activity and demonstrate that this post-translational modification serves as a critical regulatory switch for differentially modulating microtubule severing during vegetative and reproductive stages of the plant life cycle.

## RESULTS

### A triple phosphomimetic version of p60 katanin has impaired microtubule severing *in vitro*

Using the Arabidopsis Protein Phosphorylation Site Database, we identified three serine residues (S92, S147, and S199) that were experimentally found to be phosphorylated in Arabidopsis p60 katanin (KTN1; Fig. 1A). Notably, these three serine residues are highly conserved in p60 katanin subunits from other flowering plants (Suppl. Fig. S1), suggesting that they might be functionally important. To study the impact of phosphorylation of these residues on microtubule severing activity, we generated both phosphonull and phosphomimetic versions of KTN1. Single phosphonull (S92A, S147A, S199A) and phosphomimetic (S92D, S147D, S199D) modifications were created by substituting individual serine residues with either alanine or aspartate, respectively. Similarly, double and triple phosphonull and phosphomimetic mutants were generated.

**Figure 1.**
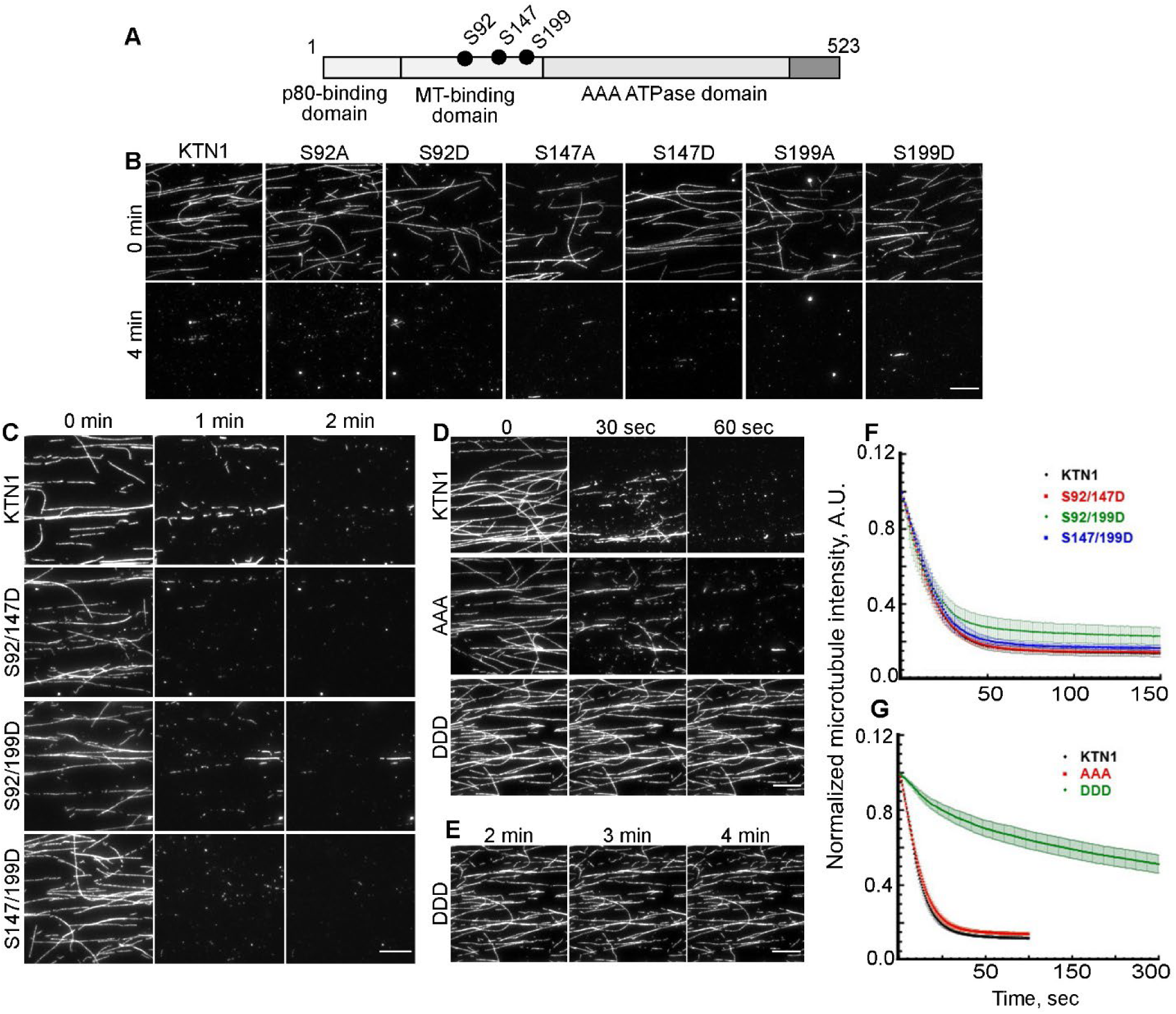
The KTN1 DDD version has decreased microtubule severing activity *in vitro*. **(A)** Diagram showing the location of the three experimentally validated serine phosphorylation sites in Arabidopsis KTN1. **(B-E)** Time course of rhodamine-labeled, taxol-stabilized microtubules incubated with 2mM ATP and 25nM of the indicated p60 katanin versions. See also Supplementary Videos 1-3. **(B)** KTN1 (n = 6), phosphomimetic and phosphonull mutants of either S92 (n = 5 each), S147 (n = 6 each), or S199 (n = 5 each). **(C)** KTN1 (n = 6) and double phosphomimetic mutations at S92 and S147 (n = 6), S92 and S199 (n = 6 movies) and S147 and S199 (n = 5). **(D)** KTN1 (n = 16), triple phosphonull (AAA, n = 15), and triple phosphomimetic (DDD, n = 13) mutant. **(E)** Extended duration of the DDD time course shown in **(D)**. Scale bar = 10 µm. (**F, G**) Plots of microtubule fluorescence signal over time from experiments in **(C)** and **(D)**. Each image in a series is normalized to the fluorescence signal of the first frame of that series. Error bars represent SEM of at least three separate protein preparations.

We used *in vitro* microtubule severing assays with taxol-stabilized, rhodamine-labeled microtubules and recombinantly purified katanin to compare the severing activity of the phosphonull and phosphomimetic mutants to wild-type KTN1. We found that all the single and double phosphonull and phosphomimetic versions of KTN1 were able to sever microtubules similar to wild type KTN1 (Figs. 1B, 1C, 1F). Next, we compared the severing activity of the triple phosphonull and phosphomimetic versions (henceforth referred to as AAA and DDD, respectively) with that of wild-type KTN1. Both KTN1 and AAA essentially eliminated the microtubule fluorescence signal within 60 sec due to extensive severing (Fig. 1D and 1G; Supplementary Videos 1 and 2). However, DDD had greatly reduced severing activity, and significant microtubule fluorescence signal persisted up to 4 minutes (Fig. 1D, 1E, and 1G; Supplementary Video 3). Additionally, higher concentrations of DDD did not restore severing activity to wild type levels (Suppl. Fig. S2). Fitting the microtubule fluorescence intensity data revealed a half-life of 16.0 s and 14.8 s for KTN1 and AAA, respectively. In contrast, DDD had a roughly 5-fold slower fluorescence decay with a half-life of 88.74 s. From these data, we conclude that the DDD modification significantly impairs the microtubule severing activity of Arabidopsis p60 katanin.

### The DDD mutant has decreased microtubule binding and ATPase activity *in vitro*

Because S92, S147, and S199 are located within the predicted microtubule-binding domain of KTN1 (Fig. 1A), modulation of microtubule binding affinity was one possible explanation for the attenuated severing activity of DDD. To test this hypothesis, we used microtubule co-sedimentation assays to measure the microtubule binding affinity of KTN1, AAA, and DDD proteins (Fig. 2A). We found that KTN1 and AAA had similar microtubule binding—both exhibited a maximum binding of about 70% and a dissociation constant of about 0.3 μM (Fig. 2B). In contrast, DDD showed a maximum binding of about 40% and a dissociation constant of about 0.7 μM (Fig. 2B).

**Figure 2.**
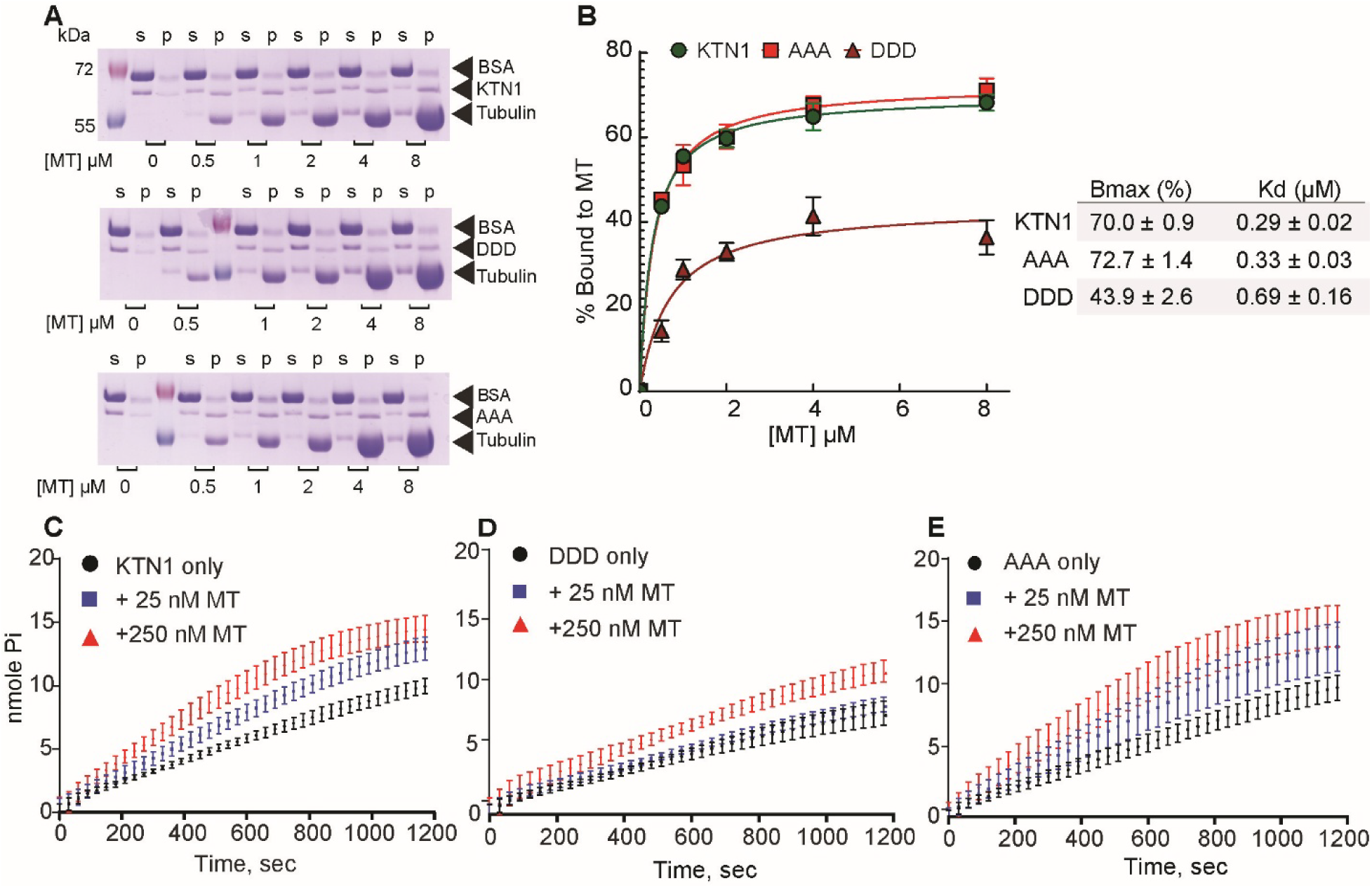
DDD has lower microtubule binding affinity and ATPase activity *in vitro*. **(A)** Representative Coomassie Blue-stained SDS-PAGE gels of microtubule co-sedimentation assays. 1 µM KTN1 (n = 3), DDD (n = 3), or AAA (n = 6) was incubated with the indicated concentration of taxol-stabilized microtubules. S, supernatant; P, pellet. Arrows identify the different protein bands. **(B)** Binding curves of KTN1, DDD, and AAA proteins corresponding to **(A)**. Each data point represents the mean ± SEM. Data were fit to a one-site saturation binding model to obtain the microtubule binding affinity and maximum amount of protein binding. **(C-E)** Plots of ATPase activity over time of 25 nM of KTN1 **(C)**, DDD **(D)**, and AAA **(E)** proteins. ATPase activity was measured as the amount of Pi release in the absence (black) or presence of either 25 nM (blue) or 250 nM microtubules (red). Each data point represents the mean ± SEM from 9 independent experiments.

Since binding to microtubules is known to stimulate the ATPase activity of katanin (Hartman et al. 1998), we next monitored the ATPase activity of KTN1, AAA, and DDD in the absence of microtubules and in the presence of 1:1 and 1:10 molar ratio of katanin to microtubules. As expected, the presence of microtubules enhanced the ATPase activity of KTN1 in a microtubule concentration-dependent manner (Fig. 2C). The AAA protein behaved like KTN1 in the absence and presence of microtubules (Fig. 2D). However, the DDD protein had about 2-fold lower basal and microtubule-stimulated ATPase activity compared to KTN1 and AAA (Fig. 2E). Together, these data indicate that the combined phosphorylation of S92, S147, and S199 negatively regulates katanin’s microtubule binding and ATPase activity.

To determine whether the AAA and DDD modifications alter interaction with the regulatory p80 subunit, we conducted yeast two-hybrid assays. Both AAA and DDD interacted with all four Arabidopsis p80 subunits with a preference for p80-1, similar to KTN1 (Suppl. Fig 3). Thus, the AAA and DDD modifications do not impact protein-protein interactions via the N-terminus of p60 katanin. In addition, these results indicate that the reduced microtubule binding and ATPase activity of DDD are not likely due to general protein misfolding.

### The DDD mutant is defective in severing interphase cortical microtubules

To determine the localization and severing activity of the katanin phosphomutants *in vivo*, we expressed GFP-tagged KTN1, AAA, and DDD in the Arabidopsis *ktn1-2* null mutant expressing an mCherry-TUB6 microtubule reporter. Live imaging of hypocotyl epidermal cells showed that both KTN1 and AAA predominantly localized as puncta at crossover and nucleation sites of cortical microtubules (Fig. 3A), in agreement with previous findings of wild type GFP-KTN1 distribution (Lindeboom et al. 2013; Zhang et al. 2013). While the DDD version also showed punctate localization along cortical microtubules, we consistently observed significant diffuse cytoplasmic signal in these plants (Fig. 3A), suggestive of reduced microtubule binding.

**Figure 3.**
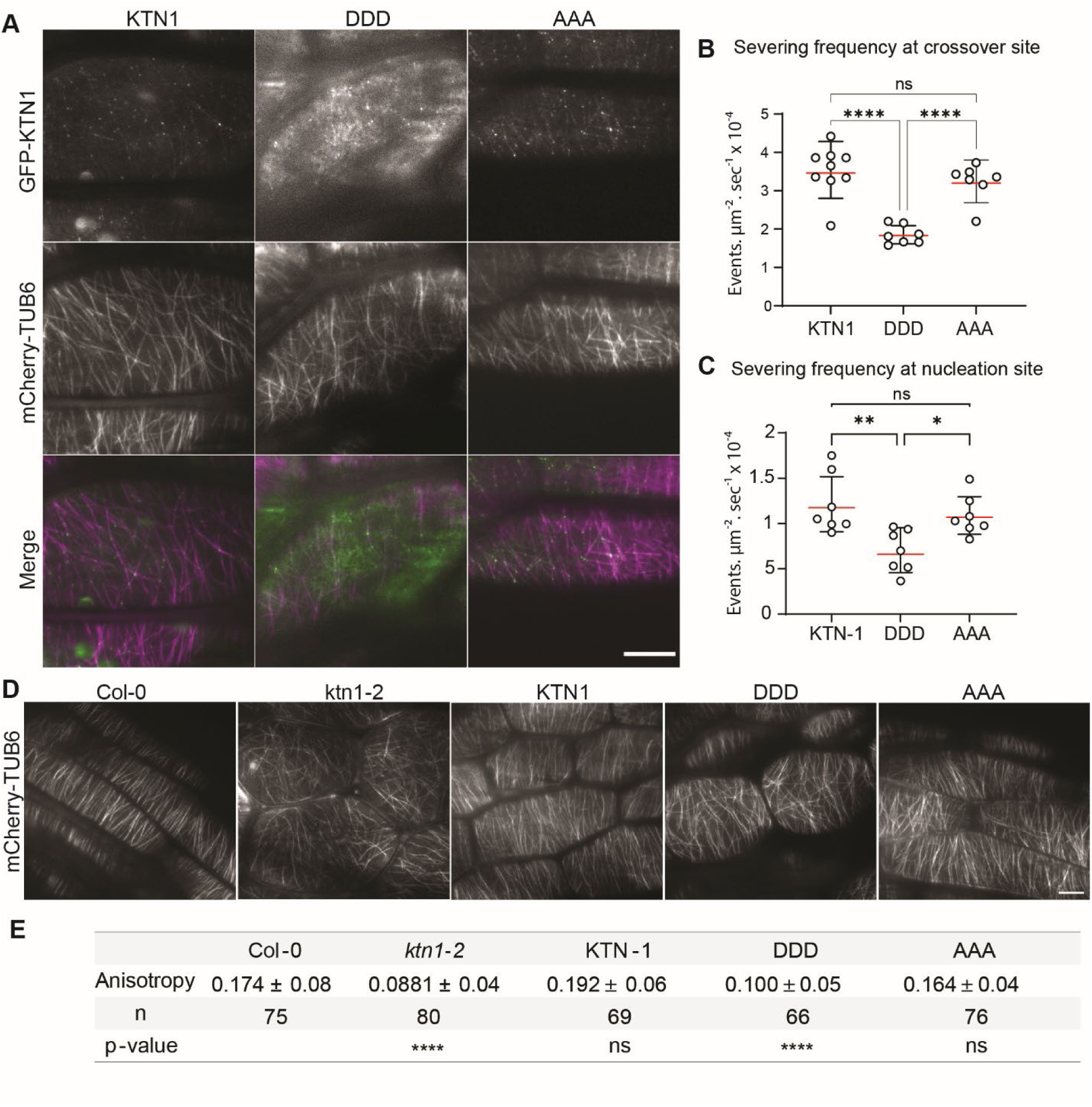
DDD is unable to fully retore cortical microtubule severing and array organization in the *ktn1-2* mutant. **(A)** Fluorescence micrographs of GFP-labeled katanin and mCherry-TUB6-labeled cortical microtubules. KTN1 and AAA localize primarily on microtubules as puncta, whereas DDD shows significant diffuse cytoplasmic signal. Scale bar = 10 µm. **(B-C)** Severing frequency of cortical microtubules at **(B)** crossover sites (n = 9 cells) and **(C)** nucleation sites (n = 7 cells) in hypocotyl epidermal cells. Red line indicates the mean and the black lines show the SD. Asterisks indicate a significant difference as determined by one-way ANOVA (ns, not significant; *P < 0.05; **P < 0.01; ****p < 0.0001). **(D)** Fluorescence micrographs of cortical microtubules in hypocotyl epidermal cells from 3-day-old seedlings. Scale bar = 10 µm. **(E)** The degree of cortical microtubule anisotropy in hypocotyl epidermal cells of the indicated genotypes. Values are mean ± SD. The number of hypocotyl cells analyzed for each genotype is shown. Asterisks indicate a significant difference as determined by one-way ANOVA (ns, not significant; ****p < 0.0001).

We next quantified the cortical microtubule severing activity of KTN1, AAA, and DDD in hypocotyl epidermal cells. Analysis of timelapse movies showed that the average severing frequency of AAA at both cortical microtubule crossover and nucleation sites were indistinguishable from KTN1 (Fig. 3B and 3C). However, the average severing frequency of DDD at both cortical microtubule locations was roughly 2-fold less than KTN1 and AAA (Fig. 3B and 3C). Consistent with the decreased severing activity of DDD, we found that *ktn1-2* plants expressing DDD had more disordered cortical microtubules compared to plants expressing either KTN1 or AAA (Fig. 3D). Quantification of cortical microtubule anisotropy using the FibrilJ tool showed that expression of KTN1 or AAA was able to restore cortical microtubule coalignment in the *ktn1-2* mutant to wild type levels (Fig. 3E). In contrast, expression of DDD only partially rescued the disorganized cortical microtubule phenotype of the *ktn1-2* mutant (Fig. 3E).

### The DDD mutant partially rescues stunted growth of the *ktn1-2* mutant

Disorganized cortical microtubule arrays in the *ktn1-2* mutant lead to reduced anisotropic cell expansion and consequently shorter roots and shoots compared to wild type plants (Bichet et al. 2001; Burk et al. 2001; Burk and Ye 2002). Since the AAA and DDD mutants differentially affected cortical microtubule organization, we wanted to characterize their impact on cell expansion and plant growth. At the seedling stage, expression of KTN1 and AAA in the *ktn1-2* mutant restored the length of light-grown roots and dark-grown hypocotyls to wild type levels (Fig. 4A-4D). In contrast, DDD was able to only partially complement the stunted growth of the *ktn1-2* seedlings (Fig. 4A-4D). We observed a similar pattern in adult plants. Both KTN1 and AAA were able to restore rosette size and inflorescence stem length to wild type levels, whereas DDD only partially complemented these morphological defects (Fig. 4E-4H).

**Figure 4.**
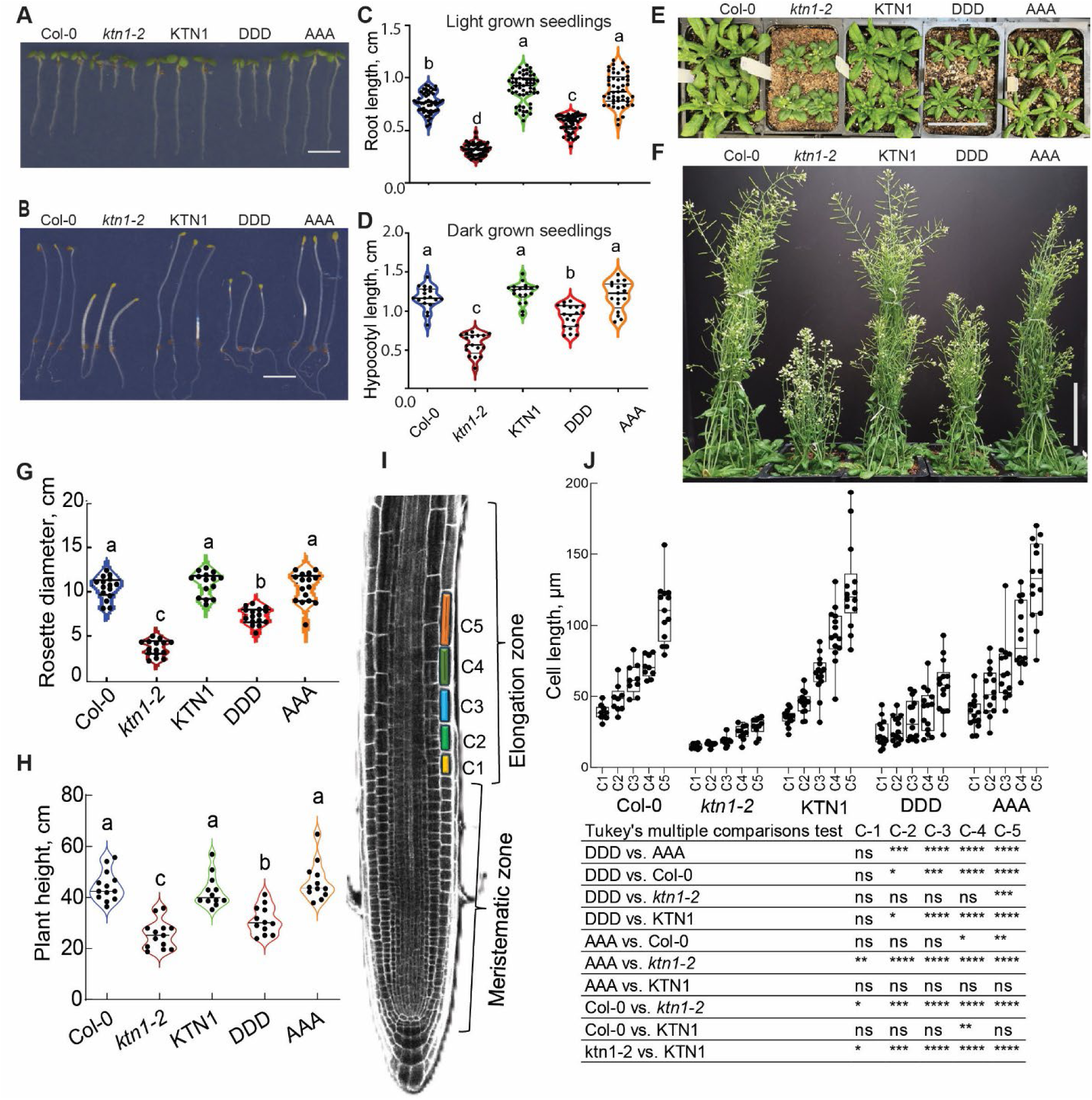
DDD only partially complements the vegetative phenotype of the *ktn1-2* mutant. **(A, B)** Images of 7-day old light-grown seedlings **(A)** and 3-day-old dark-grown seedlings **(B)** of the indicated genotypes. Scale bar = 0.2 cm. **(C, D)** Violin plots of root **(C)** and hypocotyl **(D)** length corresponding to **(A)** and **(B)**, respectively. n = 60 and 20 light-grown and dark-grown seedlings, respectively. Letters indicate statistically distinguishable groups (p < 0.05) determined by one-way ANOVA followed by Tukey’s test. **(E)** Representative images of 3-week-old soil-grown plants showing rosette size. Scale bar = 5 cm. **(F)** Representative images of 8-week-old soil-grown plants showing plant height. Scale bar = 10 cm. **(G, H)** Violin plots of rosette diameter **(G)** and plant height **(H)** corresponding to **(E)** and **(F)**, respectively. n = 15 and 12 plants, respectively. Letters indicate statistically distinguishable groups (p < 0.05) determined by one-way ANOVA followed by Tukey’s test. **(I)** Confocal micrograph of propidium iodide-stained root from a 3-day-old seedling. The first five cells of the elongation zone are colored and labeled C1-C5. **(J)** Box and whisker plots of the lengths of cells C1-C5 corresponding to **(I)**. The center line of the box plots represents the median, box limits represent the upper and lower quartiles, and whiskers represent the maximum and minimum values in the dataset (n = 10 for *ktn1-2* and Col-0; n =14 for KTN1, DDD, and AAA). The table below shows statistics determined by two-way ANOVA followed by Tukey’s multiple comparisons test (ns, not significant; *P < 0.05; **P < 0.01; ***P < 0.001; ****P < 0.0001).

To determine the effect of AAA and DDD on cell expansion, we measured the lengths of the first 5 cells in the root elongation zone of 3-day-old seedlings (Fig. 4I). In wild type plants, these cells showed a sharp increase in length while in the *ktn1-2* mutant they remained strikingly short (Fig. 4J). Consistent with the root length data, *ktn1-2* plants expressing KTN1 and AAA emulated the cell lengths of wild-type plants, whereas DDD only marginally increased cell lengths compared to the *ktn1-2* mutant (Fig. 4J).

### The AAA mutant is unable to rescue the fertility defect of the *ktn1-2* mutant

While conducting the plant experiments, we noticed that the DDD version, which was unable to fully rescue the dwarf phenotype of the *ktn1-2* mutant, unexpectedly yielded normal seed set whereas the AAA version was deficient in this function. To explain this reversal in functionality of AAA and DDD, we investigated both female and male reproductive biology.

To begin with, we used tissue clearing of mature green siliques to quantify seed filling. In Col-0, KTN1, and DDD plants, both carpels showed essentially complete seed filling, totaling about 60 seeds per silique (Fig. 5A, 5B). As reported previously for katanin null alleles (Luptovčiak et al. 2017), about 75% of *ktn1-2* siliques were empty. The remaining 25% of siliques showed only 4-8 seeds located mostly at the top or middle portion of the silique (Fig. 5A, 5B). The AAA plants had an intermediate phenotype with a mean of about 30 seeds per silique (Fig. 5A, 5B). In addition, we found that silique length was proportional to the seed set for each genotype (Fig. 5C).

**Figure 5.**
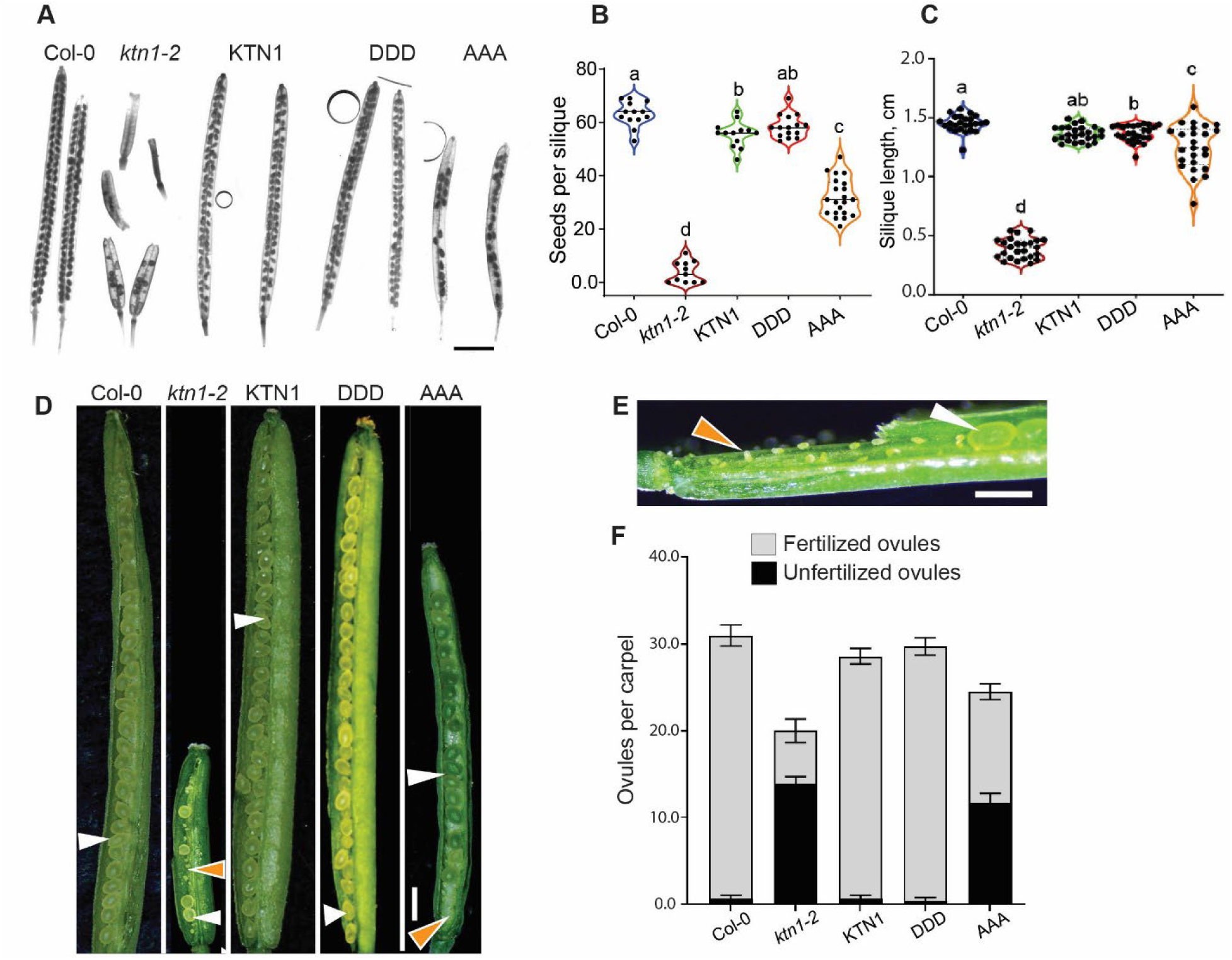
AAA only partially complements the reproductive phenotype of the *ktn1-2* mutant. **(A)** Representative image of siliques collected 10 days after anthesis and cleared with Hoyer’s reagent to visualize the total seed set. Scale bar = 2.5 mm. **(B, C)** Violin plots of total seed count per silique **(B)** and silique length **(C)**. n = 20 and 25 siliques, respectively. Letters indicate statistically distinguishable groups (p < 0.05) determined by one-way ANOVA followed by Tukey’s test. **(D)** Representative images of microdissected siliques 5 days after anthesis. Orange arrows indicate unfertilized ovules, and the white arrows indicate fertilized ovules. Scale bar = 1 mm. **(E)** Enlarged image of a silique 5 days after anthesis from a AAA plant. Orange arrow indicates an unfertilized ovule towards the base of the silique, and the white arrow indicates a fertilized ovule. Scale bar = 1 mm. **(F)** Bar graph of the number of fertilized ovules and unfertilized ovules per carpel in the indicated genotypes. Error bars represent SEM (n = 15 carpels per genotype).

To determine the cause of the low seed set in *ktn1-2* and AAA plants, we dissected carpels from each genotype at 5-7 days after anthesis and quantified the number of fertilized ovules per carpel. Consistent with the seed set data, Col-0, KTN1, and DDD plants had a mean of about 30 fertilized ovules per carpel (Fig. 5D, 5F). In contrast, both *ktn1-2* and AAA plants had fewer total number of ovules per carpel (Fig. 5D, 5F). In addition, these carpels contained a substantial number of unfertilized ovules that were primarily located towards the base of the pistil (Fig. 5D-5F). Together, our data demonstrate that the AAA mutant is unable to completely overcome defects in pollination and/or fertilization in the *ktn1-2* mutant.

### Defective pollen viability and pollen tube growth accounts for the reproductive phenotype of AAA plants

To investigate whether maternal and/or paternal tissues contribute to the reduced seed set of AAA plants, we conducted reciprocal pollination experiments. When AAA pistils were pollinated with Col-0 pollen, we observed normal pollen tube growth. At 2 hours after pollination, most Col-0 pollen tubes had grown to about 20% of the pistil length (Fig. 6B). By 24 hours after pollination, Col-0 pollen tubes had reached the base of the AAA pistil, successfully targeting essentially all ovules, similar to the behavior of Col-0 pollen on Col-0 pistils (Fig. 6A, 6C). In reciprocal experiments, AAA pollen tubes grew to a lesser extent on Col-0 pistils (Fig. 6B) and they failed to reach the base of the pistil 24 hours after pollination (Fig. 6C). In addition, the AAA pollen tubes grew erratically in the transmitting tract tissue of the ovary, often exhibiting sharp curvatures compared to relatively straight growth of Col-0 pollen tubes in AAA and Col-0 pistils (Fig. 6C and 6D). In contrast, the growth rate and trajectory of DDD pollen tubes was similar to Col-0 pollen tubes in Col-0 pistils (Fig. 6B, 6C). Based on these data, we conclude that the low fertility of AAA plants is primarily due to defective pollen function.

**Figure 6.**
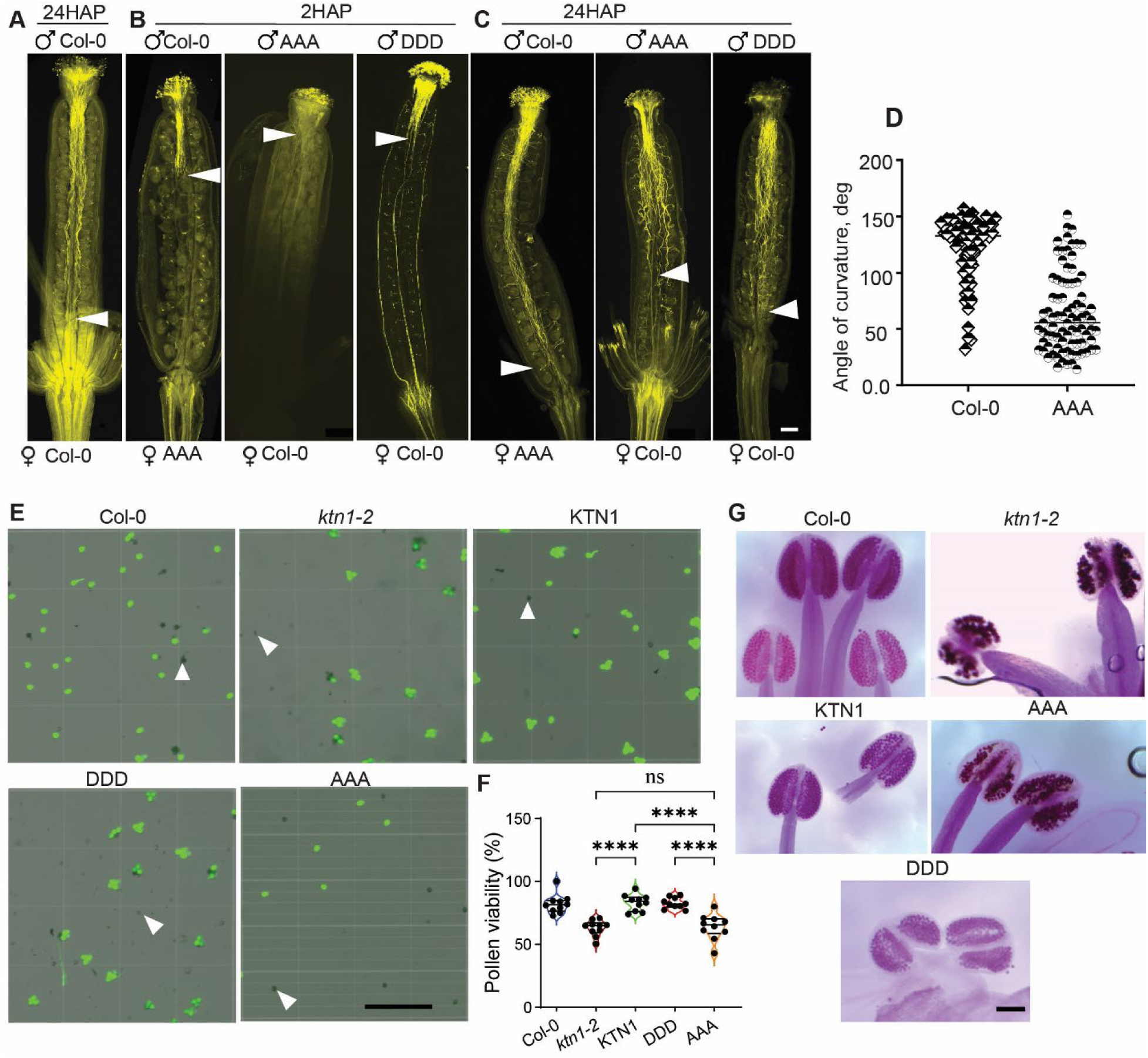
AAA pollen grains show aberrant pollen tube growth and reduced viability. **(A-C)** Representative images of aniline blue-stained pistils showing pollen tube growth. Arrowheads indicate the extent of pollen tube growth. Scale bar = 200 µm. **(D)** Angle of curvature of pollen tubes within the pistil. Deviation from straight growth was measured with respect to the proximal end of pollen tubes. Perfectly straight growth would lead to an angle of 180° while wavy growth leads to smaller angles. **(E)** Pollen viability assessed using fluorescein diacetate, which labels viable pollen. White arrows indicate dead pollen. Scale bar = 0.5 mm. **(F)** Percentage of viable pollen corresponding to **(E)**. Asterisks indicate a significant difference as determined by one-way ANOVA (ns, not significant; ****p < 0.0001). **(G)** Alexander staining of pollen grains within intact anthers. Viable pollen stain light purple while dead pollen stain dark green. Scale bar = 1 mm.

Next, we tested pollen viability by staining freshly collected pollen grains with fluorescein diacetate, which produces green fluorescence in living cells (Fig. 6E). Among all the genotypes tested, *ktn1-2* and AAA had low pollen viability whereas pollen viability of KTN1 and DDD was indistinguishable from wild type (Fig. 6F). Since *ktn1-2* pollen is known to have low pre-dehiscence viability (Luptovčiak et al. 2017), we performed Alexander staining on stage 12 flowers of all genotypes to determine if the low pollen viability of AAA was a developmental defect or a result of post-dehiscence environmental factors. We found that Col-0, KTN1, and DDD pollen grains stained uniformly purple, indicating normal development and high viability (Fig. 6G). As expected, *ktn1-2* showed greatly reduced pollen viability, evidenced by dark green staining of pollen grains within the anther (Fig. 6G). The AAA pollen exhibited an intermediate phenotype, with a mix of viable (light purple) and non-viable (dark green) pollen grains (Fig. 6G). Taken together, our data indicate that the AAA version of katanin induces male developmental defects that leads to reduced pollen viability and abnormal pollen tube growth.

### The AAA mutant leads to mitotic spindle and phragmoplast abnormalities

Katanin is known to play an important role in the correct formation of the spindle apparatus in both plants and animals (Sonbuchner et al. 2010; Loughlin et al. 2011; Panteris et al. 2011; Komis et al. 2017). In addition, loss of katanin function is associated with faulty phragmoplast and cell plate formation in plants (Panteris et al. 2011, 2021; Komis et al. 2017). To determine whether AAA and DDD impacted cell division, we analyzed the morphology of the spindle apparatus and phragmoplast in dividing cells in the meristematic zone of Arabidopsis roots. In Col-0 roots, the structure and orientation of spindles was predominantly normal, with only a minor proportion of spindles being obliquely oriented (Fig. 7A-7E). Similarly, the DDD genotype displayed normal spindles like Col-0 (Fig. 7A-7E). In contrast, the AAA genotype showed anomalies such as obliquely oriented spindles and spindles positioned at the cell periphery instead of in the center (Fig. 7A-7E). Spindles in AAA also tended to be distorted in terms of their bipolar structure (Fig. 7A) and were on average longer than in Col-0 and DDD cells (Fig. 7F). In keeping with the occurrence of spindle defects in AAA, we found that the signal of GFP-AAA at spindle poles was lower compared to the signal of GFP-KTN1 and GFP-DDD (Fig. 7G, 7I).

**Figure 7.**
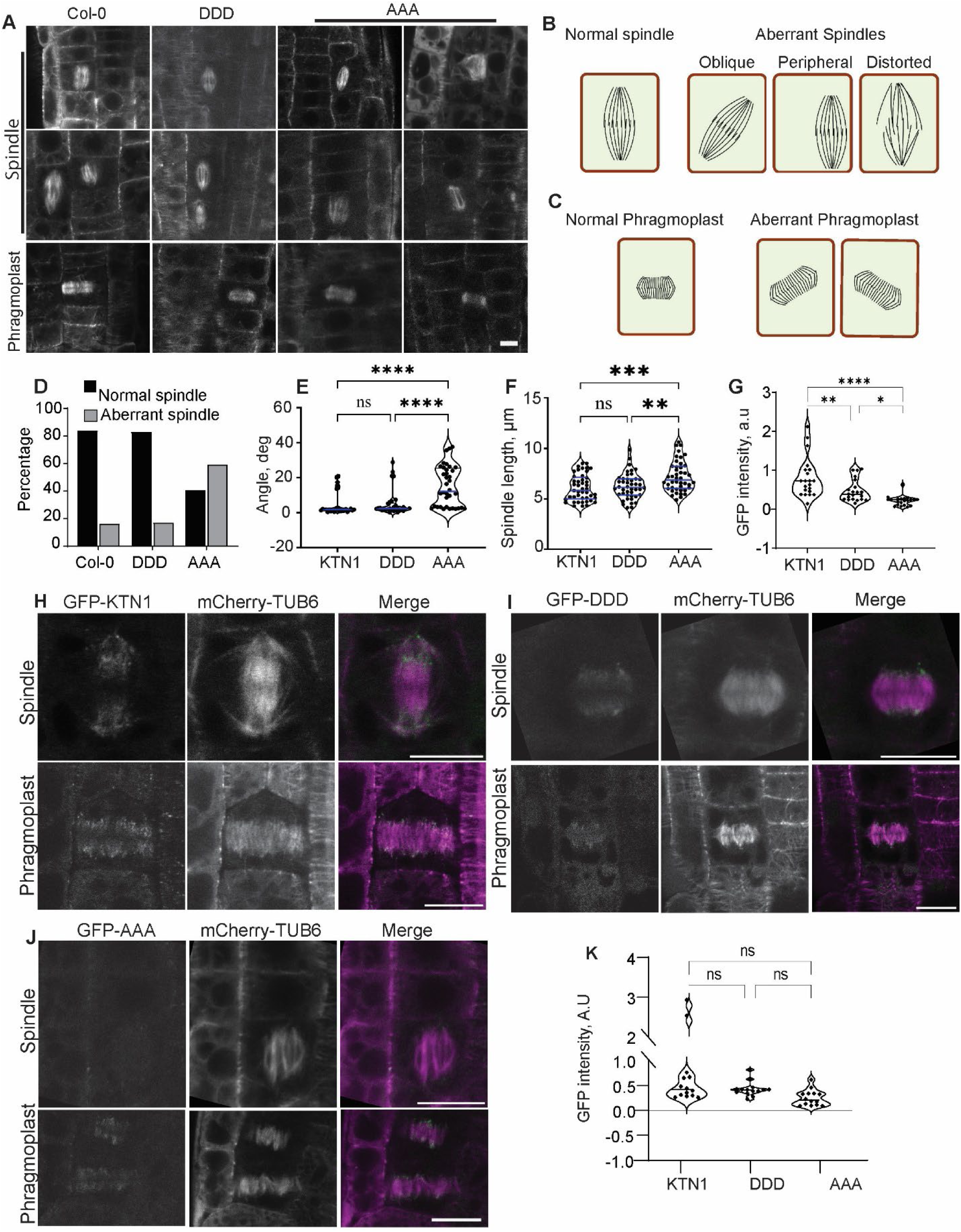
AAA plants exhibit anomalous mitotic spindles and aberrant phragmoplast orientation. **(A)** Confocal micrographs of mitotic spindles and phragmoplasts in meristematic root cells. Scale bar = 5 µm. **(B, C)** Illustration of normal and aberrant spindles **(B)** and phragmoplasts **(C)**. **(D)** Percentage of normal and anomalous spindles in the indicated genotypes (100 = spindles per genotype). **(E)** Violin plots of the angle between the long axis of the spindle and the anticlinal cell surface. N = 30 for KTN1, 29 for DDD, and 36 for AAA. **(F)** Violin plots of spindle length measured as the pole-to-pole distance. n = 45 for KTN1, 44 for DDD, and 40 for DDD. **(G)** Violin plots of GFP fluorescence intensity at spindle poles. n = 21 for KTN1, 20 for DDD, and 20 for AAA. Asterisks in **(E-G)** indicate a significant difference as determined by two-way ANOVA followed by Tukey’s multiple comparisons test (ns, not significant; *P < 0.05; **P < 0.01; ***P < 0.001; ****P < 0.0001). **(H-J)** Confocal micrographs showing the localization of GFP-KTN1 **(H)**, GFP-DDD **(I)**, and GFP-AAA **(J)** on the spindle apparatus and phragmoplast in dividing root cells expressing mCherry-TUB6. Scale bar = 5 µm. **(K)** Violin plots of GFP fluorescence intensity at the phragmoplast distal zone. n = 14 for KTN1, 14 for DDD, and 13 for AAA. Statistical significance was determined by two-way ANOVA followed by Tukey’s multiple comparisons test (ns, not significant).

In AAA roots, expanding phragmoplasts were also often obliquely oriented compared to Col-0 roots (Fig. 7A, lower panel). However, the signal intensity and distribution of GFP-AAA in phragmoplasts was similar to that of GFP-KTN1 and GFP-DDD (Fig. 7H-7K). In addition, we did not find any oblique or otherwise abnormal cell plates in AAA roots, suggesting that abnormal phragmoplast trajectories were corrected at later stages of cytokinesis.

### Meiotic spindles in pollen mother cells show severe defects in the AAA mutant

To determine whether AAA and DDD impact meiosis, we examined pollen mother cells which undergo two meiotic divisions to produce four haploid microspores. For this purpose, we isolated pollen sacs from developing anthers of stage 8-9 flowers and visualized meiosis II. Meiotic spindles in both Col-0 and DDD anthers appeared normal: they contained dense microtubules and had unfocused poles as expected for acentrosomal spindles (Fig. 8A, 8B). In striking contrast, the meiotic spindles of AAA had significantly fewer microtubules and sharply focused poles (Fig. 8A, 8B). Importantly, the microtubule signal intensity in the surrounding somatic cells was equivalent across all genotypes (Fig. 8C), demonstrating that the attenuated microtubule density is specific to meiotic cells in AAA plants.

**Figure 8.**
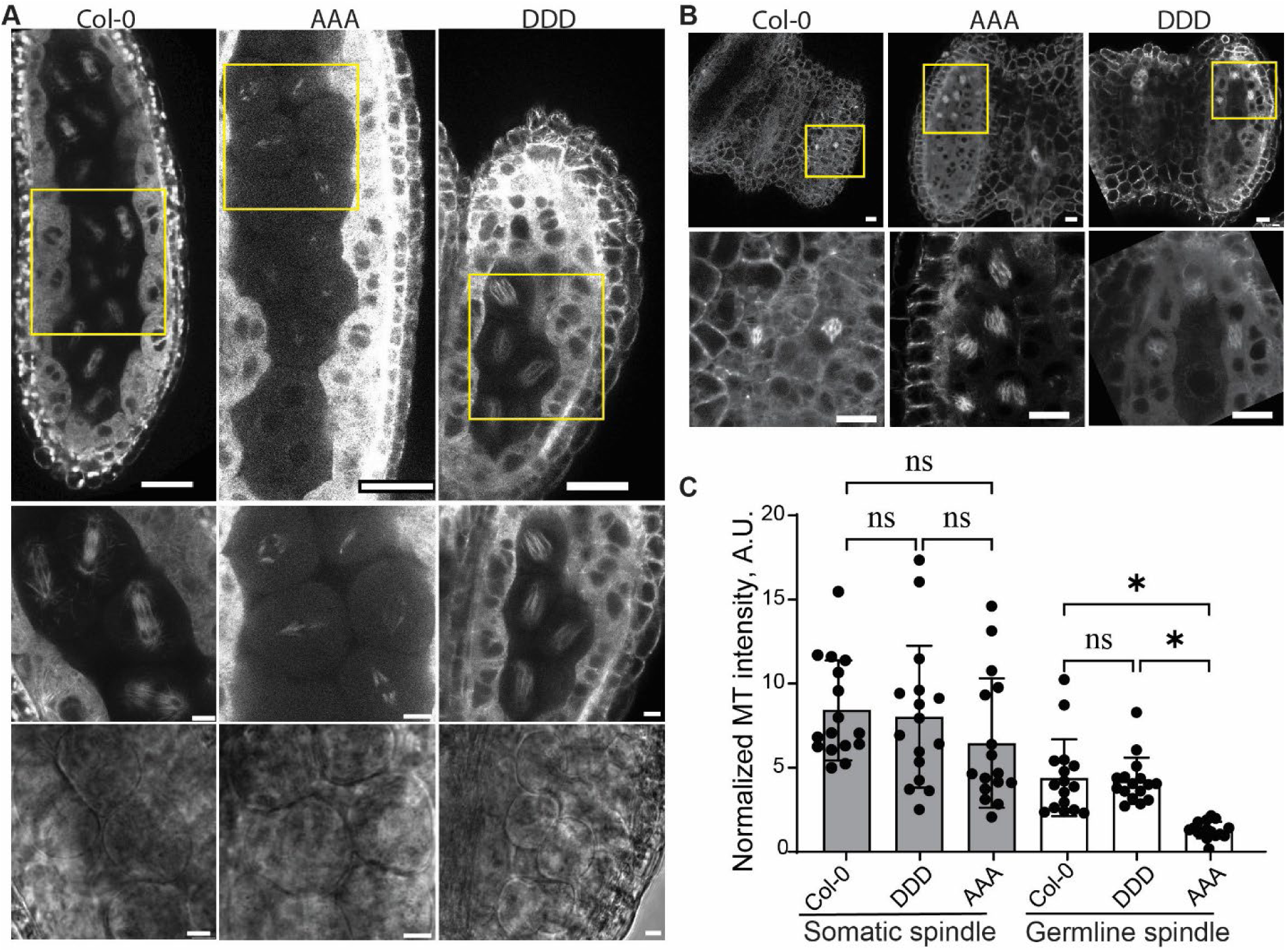
AAA expression causes severe meiotic spindle defects. (A) Confocal micrographs of anthers isolated from stage 8-9 flowers. The upper panels show meiotic spindles labeled by mCherry-TUB6. The middle panels are a closeup view of the yellow boxes in the upper panel. The lower panels show bright-field images of the middle panel. Scale bar = 10 µm. (B) Confocal micrographs of mitotic spindles in somatic cells of anthers. The lower panels are a closeup view of the yellow boxes. Scale bar = 10 µm. (C) Plots of mCherry-TUB6 fluorescence intensity per unit area of spindle apparatus from somatic and germ cells of anthers. Asterisks indicate a significant difference as determined by two-way ANOVA followed by Tukey’s multiple comparisons test (ns, not significant; *P < 0.05).

## DISCUSSION

Katanin-mediated microtubule severing plays a critical role in the construction and rearrangements of microtubule arrays throughout the plant cell cycle. However, the mechanisms by which plants regulate katanin activity to fine-tune the amount of microtubule severing at different stages of the cell cycle remains unknown. In this study, we reveal that phosphorylation at three serine residues in the N-terminal domain of Arabidopsis p60 katanin acts as a molecular switch, enabling differential control of microtubule severing during interphase-based vegetative growth and meiosis-dependent reproductive development.

Our biochemical analysis revealed that combined phosphorylation of serines 92, 147, and 199 is both necessary and sufficient to inhibit microtubule severing activity. The triple phosphomimetic DDD modification impairs both microtubule binding affinity and ATPase activity, reducing severing efficiency approximately 5-fold compared to wild-type KTN1. Importantly, higher concentrations of DDD does not restore severing activity, indicating that combined phosphorylation directly inactivates KTN1’s catalytic activity rather than modulating its oligomerization state. This mechanism contrasts with the phosphoregulation of *Xenopus* katanin at serine 131, which is thought to suppress oligomerization to tune severing activity in a concentration-dependent manner (Whitehead et al. 2013). The more stringent inhibition mechanism in Arabidopsis, requiring phosphorylation of three serine residues, may reflect the distinct cellular contexts and developmental processes that plant katanin must regulate. The conservation of these three serine residues among flowering plants suggests that this regulatory mechanism is likely an evolutionarily important solution for controlling microtubule severing in a cell cycle-dependent manner.

Despite the reduced microtubule binding affinity of DDD, our live imaging showed that DDD still localizes to cortical microtubule nucleation and crossover sites, albeit with substantial cytoplasmic signal. Furthermore, DDD exhibits normal signal intensity at spindle poles and phragmoplast distal zones during mitosis. Since our yeast two-hybrid assays demonstrated that the DDD modification does not impair binding to the regulatory p80 subunits, it is likely that DDD is recruited normally to specific microtubule sites through protein-protein interactions mediated by the p80 subunits or other targeting factors such as the Msd1-Wdr8 complex at cortical nucleation sites (Yagi et al. 2021) and CORD proteins at the phragmoplast (Sasaki et al. 2019). Therefore, we propose that this phosphoregulatory mechanism works by inhibiting catalytic activity at sites where katanin is already positioned, rather than by preventing recruitment altogether.

A striking finding from our study is that the AAA and DDD modifications have opposite effects on vegetative growth versus reproductive development, revealing fundamentally different requirements for katanin activity during interphase compared to meiotic cell divisions. Plants expressing DDD showed reduced severing at cortical microtubule nucleation and crossover sites, leading to disordered cortical microtubules, reduced anisotropic cell expansion, and overall dwarf plant stature. These phenotypes mimic the defects observed in katanin loss-of-function mutants and demonstrate that high katanin activity is essential for its function during interphase. In marked contrast, mitotic and meiotic microtubule arrays appear normal in DDD plants but are disrupted in plants expressing the AAA version. This result indicates that KTN1 must be phosphorylated to downregulate its activity during cell division. The requirement for katanin inactivation is especially critical during meiosis in microspore mother cells, where plants expressing AAA exhibited dramatically reduced microtubule density in meiotic spindles, perhaps due to excessive microtubule severing. These meiotic defects provide a plausible explanation for the reduced pollen viability in AAA plants.

Downregulating katanin activity also appears to be important for normal tip growth as evident from the erratic growth trajectory of AAA pollen tubes. The persistent deviation from straight pollen tube growth in AAA plants is reminiscent of branching and wavy growth of Arabidopsis root hairs and Norway spruce pollen treated with microtubule disrupting drugs (Bibikova et al. 1999; Anderhag et al. 2000). Based on this similarity, it is plausible that AAA disrupts the pollen microtubule cytoskeleton in a way that compromises directed tip growth.

Interestingly, AAA plants display relatively modest mitotic spindle and phragmoplast abnormalities that do not culminate in aberrant cell division planes. The limited impact on mitosis might reflect redundant regulatory mechanisms operating during this phase of the cell cycle. For example, KTN1 activity could be controlled through additional phosphorylation sites or by mitotic microtubule-associated proteins that provide phosphorylation-independent protection of spindle microtubules. Alternatively, plant mitotic spindles may inherently tolerate higher katanin activity compared to meiotic spindles.

Studies in mammals have revealed that katanin plays critical roles in male gametogenesis, regulating meiotic spindle formation, cytokinesis, and spermatid remodeling events (O’Donnell et al. 2012; Smith et al. 2012; Dunleavy et al. 2017). The severe male-specific fertility defects we observed in AAA plants mirror these findings and suggest that regulated katanin activity during male gametogenesis may be a conserved requirement across kingdoms. Why the AAA modification specifically affects male fertility but not female fertility in Arabidopsis remains to be studied. It is possible that male and female meiotic spindles differ in their sensitivity to katanin activity or in their expression of compensatory regulatory mechanisms.

Our finding that N-terminal phosphorylation differentially regulates p60 katanin in vegetative and reproductive tissues has important implications for understanding how plants coordinate cytoskeletal dynamics with developmental programs. To decipher temporal control of katanin activity, a key next step is to determine the phosphorylation state of endogenous KTN1 during different cell cycle stages and developmental contexts. In addition, the identity of the kinase or kinases that phosphorylate KTN1 at these three serine residues remains unknown. Given the cell cycle-specific effects we observed, candidate kinases include mitotic regulators such as Aurora kinases or cyclin-dependent kinases that exhibit activity peaks during cell division. It will also be important to determine whether KTN1 is specifically phosphorylated during mitosis and meiosis to inhibit its activity, or whether a counteracting phosphatase maintains KTN1 in an active, dephosphorylated state during interphase. The latter possibility is supported by recent work which identified protein phosphatase PP2A as a katanin-interacting protein that dephosphorylates p60 katanin to promote cortical microtubule organization (Ren et al. 2022).

## MATERIALS AND METHODS

### Plant material and growth

All *Arabidopsis thaliana* (L.) plants used in this study are Columbia-0 (Col-0) ecotype. For growth on plates, seeds were surface sterilized using 5% (v/v) bleach for 10 min, followed by five thorough rinses with autoclaved Milli-Q water. Sterilized seeds were plated on ½-strength MS medium with Gamborg’s vitamins (Caisson Labs) supplemented with 0.3% (w/v) sucrose and 1% (w/v) phytoblend agar (Caisson Labs) and stratified for 3 days at 4°C. Plates were then moved to growth chambers with 16/8-h light/dark photoperiod of 120µmol m^-2^ sec^-1^ light intensity, 75% humidity and 24°C. For soil growth, seeds were sown in BM6 peat-and-perlite growing medium, stratified for 3 days at 4°C, and grown under continuous light at 120-140 µmol intensity, 70% humidity, and 22°C.

### Plasmid construction

For plant transformation, genomic DNA fragments encoding the AAA or DDD phosphomutant variants were synthesized *de novo* and used to replace the corresponding wild-type genomic fragment in the pBIN19-GFP-KTN1 vector (a gift from Prof. David Ehrhardt), generating KTN1 promoter-driven GFP-tagged AAA and DDD constructs.

For protein expression, site-directed mutagenesis was used to generate single, double, and triple phosphonull and phosphomimetic variants of KTN1. Mutagenic primers (Supplementary Table 1) were designed with the target substitution at the center, flanked by homologous sequence from the pGEM-KTN1 template generated before (Burkart and Dixit 2019). PCR amplification was performed using a high-fidelity polymerase, followed by a DpnI digestion to selectively degrade the methylated parental DNA template. The resulting nicked plasmid DNA was transformed into competent *E. coli*. Successful mutagenesis was initially confirmed by diagnostic restriction digest screening, exploiting either introduced or eliminated restriction sites. Sequential rounds of this process, using unique restriction sites in the pGEM-KTN1 vector for subcloning, were employed to combine mutations to create the AA/DD and AAA/DDD variant constructs. All constructs were subsequently verified by sequencing.

### Protein expression and purification

The pHMT-KTN1, pHMT-AAA, and pHMT-DDD plasmids were transformed into BL21-CodonPlus(DE3)-RIPL competent cells (Agilent Technologies). Proteins were induced and affinity-purified using Ni-NTA agarose (Qiagen) as described previously (Burkart and Dixit 2019).

### Microtubule-severing assays

Imaging chambers were constructed on microscope slides with silanized coverslips and double-sided tape (Dixit and Ross 2010). The coverslip surface was functionalized with 0.8% anti-tubulin antibody (Sigma #T4026) in SBII Buffer (20 mM HEPES, 3 mM MgCl2, 10% sucrose, pH 7.0) for 5 min and the surface then blocked with 5% pluronic F-127 (Sigma #P2443) in SBII. Subsequently, 20 µL of rhodamine-labeled porcine microtubules (a 1:100 dilution of 50 µM stock of 1:25 rhodamine-labeled porcine microtubules in SBII containing 20 µM taxol) was flowed in and incubated for 5 min, followed by two washes with SBII containing 20 µM taxol to remove unbound microtubules. The severing mix (25 nM of KTN1, AAA, or DDD, 2 mM ATP, 50 mM DTT, 800 μg/ml glucose oxidase, 175 μg/ml catalase, 22.5 mg/ml glucose, and 20 μM taxol in SBII) was then flowed in and the slide was immediately imaged by total internal reflection fluorescence microscopy. Acquisition used a 561 nm laser at 2 mW, a TRITC filter, a 150 ms exposure, and an intensification setting of 150. Images were captured at 1 s intervals for 3-5 min using a back-illuminated electron-multiplying CCD camera (ImageEM; Hamamatsu, Bridgewater, NJ) and a 582–636 nm emission filter set. Image analysis was performed using the Fiji ImageJ package (Schindelin et al. 2012).

### Microtubule co-sedimentation assays

Binding of katanin to microtubules was examined by coincubating 1 μM of either KTN1, AAA, or DDD with different concentrations of taxol-stabilized microtubules in SBII buffer supplemented with 50 μM DTT, 40 μM taxol, 0.1 mg/ml bovine serum albumin, and 2 mM AMPPNP. After incubation at room temperature for 30 min, the binding reactions were centrifuged at 39,000g for 25 min at 15°C. The supernatant and pellet fractions were run out on SDS–PAGE and stained with Coomassie Blue. Band densitometry was performed using Fiji software (Schindelin et al. 2012). To account for a small fraction of katanin pelleting in the absence of microtubules, the bound fraction at each microtubule concentration was adjusted by subtracting the amount of katanin in the pellet fraction of the no-microtubule control. Data were fit to a one-site saturation binding model, Y = B_max_*X / (K_d_ + X), in GraphPad Prism.

### ATPase assays

The ATPase activity of 25 nM of KTN1, DDD, or AAA in the presence of either 25nM or 250nM of taxol-stabilized microtubules was evaluated using a commercially available ATPase ELIPA Kit (Cytoskeleton #BK051) according to the manufacturer’s instructions.

### Transgenic plant lines

GFP-tagged KTN1, AAA, and DDD were introduced into Col-0 plants using Agrobacterium-mediated floral dip transformation (Clough and Bent 1998). Homozygous lines expressing these transgenes were then crossed with the *ktn1-2* mutant expressing mCherry fused to β-tubulin 6 (mCherry-TUB6) under the control of the Arabidopsis ubiquitin-10 promoter. Double marker lines were selected using Basta (10 mg/ml) and Hygromycin (30 mg/ml) and verified using PCR-based genotyping (see Supplementary Table 1 for primer sequences) to obtain homozygous lines.

### Plant phenotyping

For root and hypocotyl measurements, seedlings were grown vertically on a 1% agar plate for 4 days in light and dark conditions. Images of seedlings were captured using a CanoScan 4400F scanner at 600 dpi. Images of rosette leaves and inflorescence stems were captured using 3-week-old and 8-week-old soil-grown plants, respectively. Silique length was determined using the 15^th^ silique from the first open flower. Root length, hypocotyl length, rosette diameter and silique length were measured using the Fiji ImageJ package (Schindelin et al. 2012).

### Root cell length measurements

To measure the lengths of the first five epidermal cells in the root elongation zone, 3-day-old seedlings were mounted in 50 µL of 10 µg/mL propidium iodide solution to stain the cell walls. Images were collected using a 40x oil-immersion objective (NA 1.25) on a Leica SP8 confocal microscope. The dye was excited with a 552 nm laser and its emission was detected within the 562–725 nm range.

### Silique and ovule characterization

Chloral hydrate-mediated tissue clearing was employed to quantify the seed content of intact siliques. Siliques were harvested and immersed in 100% methanol overnight to fix the tissue and remove chlorophyll. Following decolorization, the siliques were sequentially rehydrated in decreasing ethanol concentrations (90%, 70%, 50%, 30%, 10% v/v) for 10 minutes each, ensuring complete submersion of the specimens. Subsequently, the siliques were placed on a glass slide and mounted with 100µl of Hoyer’s solution (chloral hydrate, water, and glycerol in an 8:2:1 v/v ratio) and incubated for 2 days. Cleared siliques were observed under a stereo dissecting light microscope (M205 FCA, Leica) equipped with a monochrome camera (DFC9000 GT, Leica). To assess the fertilization status of ovules, young siliques were incised along both sides of the replum using an insulin needle (27-gauge, 1/2-inch), and one valve was removed with fine forceps to expose the ovules. Images were captured using a stereo microscope (IVESTA3, Leica) fitted with a color camera (Flexacam c5, Leica).

### Pollen viability assays

To examine pollen viability, 20 freshly opened flowers were collected in 1.7 mL tubes containing 750 µL of pollen germination medium (PGM; composed of 18% (w/v) sucrose, 2 mM CaCl, 2 mM Ca(NO), 0.5 mM H BO, 1 mM MgSO, and 1 mM KCl, pH 7.05-7.1).

Each tube was vortexed thoroughly for at least 1 minute to suspend all flowers in PGM, followed by centrifugation at 10,000g for 5 minutes to collect the pollen grains as a yellow pellet. The supernatant, including floral debris, was carefully removed without disturbing the pellet. Fresh PGM (100 µL) was added to the pellet, and the tube was gently flicked to resuspend the pollen grains. 1 µL of fluorescein diacetate (2 mg/mL in acetone) was then added and pollen viability assessed by counting the number of stained and unstained pollen using a hemocytometer counting chamber under a stereo dissecting light microscope (M205 FCA, Leica).

For Alexander staining, 2-4 stage 12 flowers were cut at the base to remove the sepals and petals. They were placed on a microscope slide with 100 µL of Alexander staining solution (Alexander 1969; Hedhly et al. 2018). The slide was warmed briefly over a Bunsen burner for 10 seconds, with additional staining solution added as needed. A coverslip was placed and sealed with nail polish to prevent drying. The slides were incubated overnight at 40°C and observed under a stereo microscope with apochromatic optics (Leica S8APO) equipped with a color camera (Leica EC3). Viable and nonviable pollen were indicated by purple and green color, respectively.

### Pollination experiments

For reciprocal pollination experiments, 8-10 unopened flowers were emasculated on the morning of day 1. On day 2, these emasculated flowers were hand-pollinated with pollen isolated from freshly opened flowers of either Col-0, AAA, or DDD plants. At 2 and 24 hours after pollination, the pistils were collected and fixed in Carnoy’s buffer (ethanol, chloroform, and glacial acetic acid in a 6:3:1 v/v ratio) for 2-3 hours under vacuum. Following fixation, the pistils were sequentially rehydrated with 70% ethanol for 10 minutes, followed by 50%, 20%, and 10% ethanol (v/v), each for 10 minutes, and then incubated in 8 M NaOH overnight. The pistils were subsequently rinsed with distilled water to remove residual NaOH and incubated in decolorized aniline blue staining solution for at least 30 min at room temperature in the dark (Lu et al. 2011). After staining, a coverslip was placed over the pistils with a drop of glycerol, and they were observed under a Leica SP8 confocal microscope equipped with an HC-PL APO 10x objective (NA 0.4). Aniline blue was excited with a 405 nm laser, and emission was detected in the 400-519 nm range. Images were captured in tile mode and stitched with a 10% overlap. Pollen tube growth trajectories were measured in the transmitting tract of the upper portion of the ovary.

### Mitotic and meiotic spindle morphology

Mitotic microtubule structures were visualized in the root meristematic zone from 3-day-old seedlings of Col-0, GFP-KTN1, GFP-AAA, and GFP-DDD expressing mCherry-TUB6. For live imaging of meiotic spindles, flower buds at stages 8-9 from adult plants were carefully dissected under a stereomicroscope (IVESTA3, Leica) to isolate intact anthers. The isolated anthers were immediately placed in a small drop of ½-strength MS medium on glass slides prepared with double-sided tape, and a coverslip was gently placed over the specimens. Fluorescence micrographs were captured using a Leica SP8 confocal microscope equipped with a 63x oil-immersion objective (NA 1.4). GFP and mCherry were excited by 488 nm and 561 nm lasers, respectively, and their emissions detected at 500-545 nm and 580-630 nm, respectively.

Measurements of spindle and phragmoplast angles with respect to the root long axis and spindle microtubule fluorescence intensity were performed using Fiji.

### Statistical analysis

For statistical comparisons, unpaired two-tailed t-test was used to compare two unrelated datasets and one-way analysis of variance (ANOVA) plus post-hoc Tukey test was used to compare more than two unrelated datasets, unless stated otherwise in the figure legend. Statistics and graph generation were performed using either GraphPad Prism (version 10.1.0) or Excel softwares. Exact statistical tests, n numbers, and P value cut-offs are provided in figure legends.

## ACKNOWLEDGEMENTS

This work was supported by the National Institute of General Medical Sciences of the National Institutes of Health under award number R35GM139552 (R.D.).

## AUTHOR CONTRIBUTIONS

R.D., G.B., and V.A. conceived the study and designed the experiments. G.B. conducted the in vitro microtubule severing, microtubule co-sedimentation, and yeast two-hybrid experiments.

G.B. and R.B. generated plasmid constructs and transgenic plant lines. V.A. conducted all the plant phenotyping, pollination, and live imaging experiments and analyzed the data. V.A. and R.D. wrote the original draft, and V.A., G.B., R.B., and R.D. reviewed and edited the paper. R.D. acquired funding and supervised the project.

## COMPETING INTERESTS

The authors declare no competing interests.

## Notes

### Competing Interest Statement

The authors have declared no competing interest.

