## Supplemental figures and table for "N-terminal phosphorylation inhibits Arabidopsis katanin and affects vegetative and reproductive development in opposite ways"

```
P.patens       KWNKCKKDLMEEFNIVKQIDTERQAFRDAPEIYVSHPPN--------------R--SDSS
T.aestivum     KWMNCKKAISEEVEIVRQLDAQLKSFKEAPGTMRSSSPPIRSNNKSFLFQPLDEYPTSSP
O.sativa       KWLNCKKAICDEVEIVKQLDAQLKAFKEAPGARRSSSPPIRSNNKSFVFQPLDEYPTSSP
Z.mays         KWMNCKKAISEEVEIVKQLDAQLKSLKEAPGTRRSSSPPIRS-NKSFVFQPLDEYPTSSP
S.bicolor      KWMNCKKAISEEVEIVKQLDAQLKSLKEAPGTRRSSSPPIRSNNKSFVFQPLDEYPTSSP
A.thaliana     KWMNVKKAIMEETEVVKQLDAERRAFKEAPTGRRAASPPINTK-SSFVFQPLDEYPTSSG
C.sativa       KWMNVKKAIMEETEVVKQLDAERRAFKDAPTGRRAASPPINTK-SSFVFQPLDEYPTSS-
C.rubella      KWMNVKKAIMEETEVVKQLDAERRAFKEAPTGRRAASPPINTK-SSFVFQPLDEYPTSS-
S.lycopersicum KWMNVKKAISEETEVVKQLDAEKRSFKEVPMGRRPNSPPISTK-SSFVFQPLDEYPTSSG
P.vulgaris     KWMNVKKALSEETEVVKQLDAERRAFKENPIGRRAVSPPISAKSSSFVFQPLDEYPTSSS
M.truncatula   KWMNVKKALSEETEVVKQLDADKRAFKDNPIGRRPPSPPISTK-SSFVFQPLDEYPTSSN
P.trichocarpa  KWMNLKKSLSEETEIVKQLDADRRAFKEAPAARRVASPPIHAK-SSFVFQPLDEYPTSSA
G.hirsutum     KWMNVKKALSEEAEVVKQLDAERRSFKEAPNGRRPSSPPIHAK-SSFVFQPLDEYPTSSG
V.vinifera     KWMNVKKALSEETEVVKQLDAERRAFKEIPGGRRPSSPPISTK-SSFVFQPLDEYPTSSG

P.patens       RSDPNDDPDVWRPPSRDIPSQYGTRRPTKAGQMAAAAQAKRMAAYARGAGDRTASQVGRG
T.aestivum     --STFDDPDVWAPPRDTNTP----NRRSARGQSSAR-KSSQDGAWARGPSKTGTPSRGAK
O.sativa       --APFDDPDVWAPPRD--TP----NRRPTRGQSSTR-KSSQDGAWARGPSKTGTPSRGAK
Z.mays         --APFDDPDVWAPPRD--TP----TRRPTRGQSSAR-KSSQDGAWARGSSRTGTPSRSAK
S.bicolor      --APFDDPDVWAPPRD--TP----TRRPARGQSSAR-KSSQDGAWARGSARTGTPSRSTK
A.thaliana     G-GPMDDPDVWRPPTRDVTS----RRPARAGQTGTR-KSPQDGAWARGPTTRTGPASRGG
C.sativa       G-GPMDDPDVWRPPTRDVTS----RRPARSGQSGMR-KSPQDGAWARGPTTRTGPASRGG
C.rubella      G-GPMDDPDVWRPPTRDVTT----RRPARSAQSGMR-KSPQDGAWARAPATRTGPASRGG
S.lycopersicum --APMDDPDVWRPPSRDT-T----RRPGRAGQGGMR-KSPQDGAWARASTTRTGTTIRGG
P.vulgaris     GPGPVDDPDVWRPPSRDT-S----RRPARSGQVGTR-KSPQDGAWARGATARSG-STGRG
M.truncatula   --SSFDDPDVWRPPSRDT-S----RRPGRGGQVGTR-KSNQDSKWARGATTKTTSTTGRG
P.trichocarpa  --APIDDPDVWRPPSRDTTS----RRPTRASQVGLR-KSPQDGAWARGASTRTG-TTGRG
G.hirsutum     --APMDDPDVWRPPSRDTST----RRPARGGQAGMR-KSPQDGISGRGS-TRTA-ATGRG
V.vinifera     --APMDDPDVWRPPSRDTTS----RRSARSGQVGMR-KTSQDGTWPRGS-TRGG-TAPRG

P.patens       NKVTPSASSKTIGQGA-GMGAKGSSVSGTTVPRKSTSSTGRSTPSKRESLNGAPEEEVPR
T.aestivum     PSG----SKASSAVRSSTASSTGG--------RKGK-S----SSSKADSASSDAEE----
O.sativa       PNG----SKGNSGARSSTASSTGG--------RKGK-S----SSSKADPVSSDAEE----
Z.mays         PNG----IKGGAVK--STASNSSV--------RKGK-Q----SSNKADSTSSDAEE----
S.bicolor      PNG----SKGGSTVKSSAASNSSV--------RKGK-P----SSSKADSASSDAEE----
A.thaliana     --RGGATSKSTAGARSSTAGKKGA--------ASK--------SNKAESMNGDAED----
C.sativa       --RGGATSKSTAGARSSTAGKKGA--------ASK--------STKADSTNGDAEE----
C.rubella      --RGGATSKSAAGSRSSAAGKKGA--------ASK--------STKADSTNGDAED----
S.lycopersicum RTSG--STKAGTGVRSSTTGKKAP--------GKS---A------KSDSTDGEPEE----
P.vulgaris     GAKGGAISRVNSGTRASATGKKGN--------VSGK-STKTDTAAVTNGQNGDSED----
M.truncatula   AKAGGATSRTNSGARGSTPVKKGG--------ASGK-SSNTD------AVNGDGED----
P.trichocarpa  AKT-AASSRVNSGVRASTTGKKGT--------GTGK-SGRGD------SANGDAED----
G.hirsutum     AKA-GASSRTNTGVRGSTTGKKGT--------GSGK-PSKGD------SANGDAED----
V.vinifera     GKS-GASSRTHSGVRASTTGRKGS--------SKSN-TDKAD------SANGDAED----
```

**Supplementary Figure S1.** Amino-acid sequence alignment of p60 katanin from various plants species. Arrows point to Arabidopsis serine residues that are phosphorylated and the focus of this study.

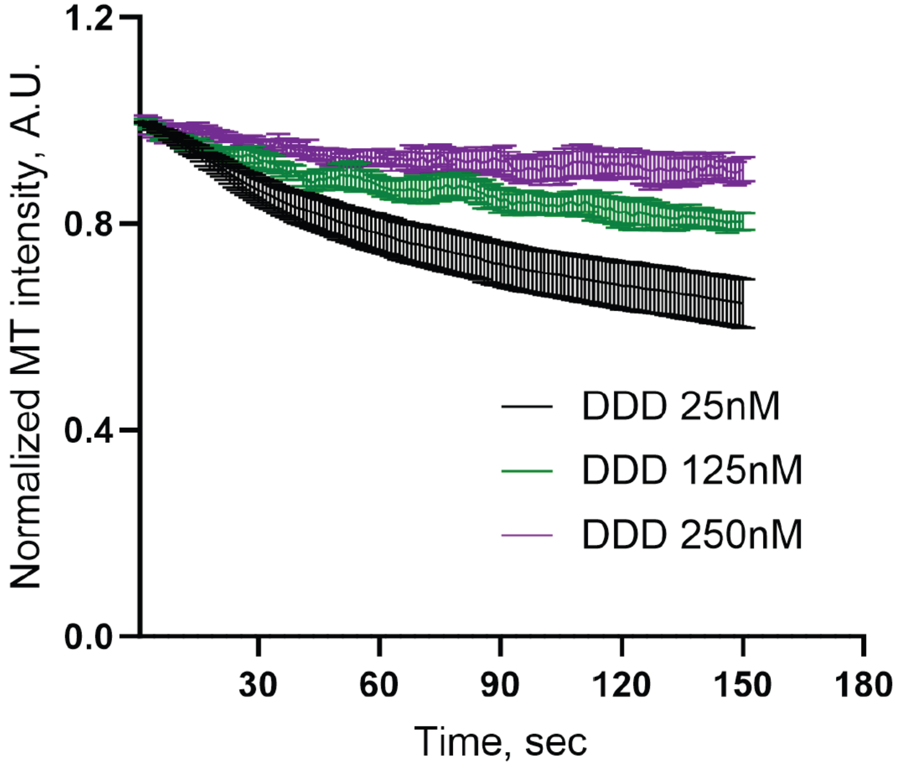


**Supplementary Figure S2.** Effect of increasing concentrations of DDD on microtubule severing activity. Plot of microtubule fluorescence signal over time from time courses of rhodamine-labeled, taxol-stabilized microtubules in the presence of 2 mM ATP and either 25 nM, 125 nM, or 250 nM of DDD. Each image in a series is normalized to the fluorescence signal of the first frame of that series. Error bars represent SD.

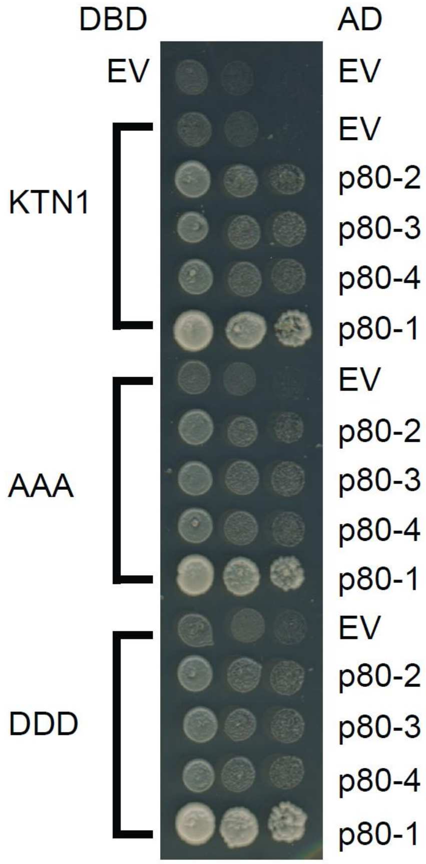


**Supplementary Figure S3.** Directed yeast-2-hybrid assay examining interaction of KTN1, AAA, and DDD with the four p80 katanin isoforms of Arabidopsis. DBD, DNA binding domain. AD, activation domain. EV, empty vector.

**Supplementary Table 1**. Sequences of primer used in this study.

| Primer Name | Primer sequence | Purpose |
|---|---|---|
| ktn1-2 Geno-R | GCAGATCCAAACTCAGAGAGC | Genotyping ktn1-2 locus |
| SAIL_LB | GAATTTCATAACCAATCTCGATACAC | T-DNA primer |
| ktn1-2 Geno-F | GACTCTCTCTGCAATTCTCGG | Genotyping ktn1-2 locus |
| EGFP Int F | GTGCAGTGCTTCAGCCGCT | Transgene detection |
| KTN1 Int R | GCTTGTTGATCTGAGCAATGGC | Transgene detection |
| ktn1-2 Geno-R2 | GACTCTCTCTGCAATTCTCGG | Sequencing KTN1 genomic phosphomutants |
| KTN1 phospho-F | GGAAGAGACGGAAGTTGTGAAGC | Sequencing KTN1 genomic phosphomutants |
| p60 AscI S92A F | GCTCCCACTGGGCGGCGCGCCGCT GCTCCACCTATCAATACC | Phosphonull mutation of S92 |
| p60 AscI S92A R | GGTATTGATAGGTGGAGCAGCGGC GCGCCGCCCAGTGGGAGC | Phosphonull mutation of S92 |
| p60 AscI S92D F | GCTCCCACTGGGCGGCGCGCCGCT GATCCACCTATCAATACC | Phosphomimetic mutation of S92 |
| p60 AscI S92D R | GGT ATT GAT AGG TGG ATC AGC GGC GCG CCG CCC AGT GGG AGC | Phosphomimetic mutation of S92 |
| p60 MluI S147A F | GCTGGTCAAACTGGTACGCGTAAAGCA CCTCAAGATGGGGCT | Phosphonull mutation of S147 |
| p60 MluI S147A R | AGCCCCATCTTGAGGTGCTTTACGCGT ACCAGTTTGACCAGC | Phosphonull mutation of S147 |
| p60 MluI S147D F | GCTGGTCAAACTGGTACGCGTAAAGAC CCTCAAGATGGGGCT | Phosphomimetic mutation of S147 |
| p60 MluI S147D R | AGCCCCATCTTGAGGGTCTTTACGCGT ACCAGTTTGACCAGC | Phosphomimetic mutation of S147 |
| p60 EciI S199A F | CTCAAAATCTAACAAGGCGGAGGCTAT GAATGGTGATGC | Phosphonull mutation of S199 |
| p60 EciI S199A F | GCATCACCATTCATAGCCTCCGCCTTG TTAGATTTTGAG | Phosphonull mutation of S199 |
| p60 EciI S199D F | CTCAAAATCTAACAAGGCGGAGGATAT GAATGGTGATGC | Phosphomimetic mutation of S199 |
| p60 EciI S199D F | GCATCACCATTCATATCCTCCGCCTTG TTAGATTTTGAG | Phosphomimetic mutation of S199 |

**Supplemental Video 1**. In vitro microtubule severing assay with taxol-stabilized, rhodamine-labeled microtubules incubated with 25 nM KTN1 and 2mM ATP.

**Supplemental Video 2**. In vitro microtubule severing assay with taxol-stabilized, rhodamine-labeled microtubules incubated with 25 nM AAA and 2mM ATP.

**Supplemental Video 3**. In vitro microtubule severing assay with taxol-stabilized, rhodamine-labeled microtubules incubated with 25 nM DDD and 2mM ATP.
